# Eye-movement reinstatement and neural reactivation during mental imagery

**DOI:** 10.1101/107953

**Authors:** Michael B. Bone, Marie St-Laurent, Christa Dang, Douglas A. McQuiggan, Jennifer D. Ryan, Bradley R. Buchsbaum

## Abstract

Half a century ago, Donald Hebb posited that mental imagery is a constructive process that emulates perception. Specifically, Hebb claimed that visual imagery results from the reactivation of neural activity associated with viewing images. He also argued that neural reactivation and imagery benefit from the re-enactment of eye movement patterns that first occurred at viewing (fixation reinstatement). To investigate these claims, we applied multivariate pattern analyses to functional MRI (fMRI) and eye-tracking data collected while healthy human participants repeatedly viewed and visualized complex images. We observed that the specificity of neural reactivation correlated positively with vivid imagery and with memory for stimulus image details. Moreover, neural reactivation correlated positively with fixation reinstatement, meaning that image-specific eye movements accompanied image-specific patterns of brain activity during visualization. These findings support the conception of mental imagery as a simulation of perception, and provide evidence of the supportive role of eye-movement in neural reactivation.

---

The idea that mental imagery involves the reactivation of neural activity patterns elicited at perception has now been firmly established^1–7^. To date, much of the work on the neural basis of visual imagery has examined the phenomenon as if mental images were visual snapshots appearing in their totality to a passive inner observer, with few exceptions^8^. However, mental imagery is an active, constructive process^9,10^ that is subject to the very kinds of capacity limitations that constrain perception and working memory^11^, leading some to propose that people engage with mental images in much the same way as they explore the sensory world—using eye-movements to shift the focus of attention to different parts of a mental image^12–19^. To date, however, there is scant neuroscientific evidence showing that eye-movement patterns are related to the neural representations that support mental imagery for complex visual scenes.

In a seminal paper^12^, Donald O. Hebb proposed a theory of mental imagery comprising three core claims: 1) imagery results from the reactivation of neural activity associated with the sequential perception of “part-images” (i.e. the spatially organized elements of a mental image); 2) analogous to the role of saccades and fixations during perception, eye movements during imagery temporally organize the neural reinstatement of such “part-images”, thereby facilitating imagery by reducing interference between different image parts; and, 3) the vividness and detail of mental imagery is dependent on the order (first-, second-, third-order, etc.) of neuronal cell assemblies undergoing reactivation, such that reactivation extending into lower order visual regions would elicit greater subjective vividness than reactivation limited to higher-order areas.

Hebb’s first claim that imagery requires the reinstatement of perceptual neural activity has received considerable empirical support over the last decade. The advent of multi-voxel pattern analysis^20^ (MVPA) has facilitated the assessment of neural reactivation, which is when stimulus-specific activity patterns elicited at perception are reactivated during retrieval^21,22^. Researchers have consistently reported substantial similarities between neural regions activated by visual imagery and visual perception^2,3,23^, and there is now significant evidence that measures of neural reinstatement reflect the content^4–7,24^ and vividness^25–28^ of mental imagery.

Hebb’s third claim that vivid mental imagery is the result of neural reinstatement within early visual areas (e.g. V1) has also received some neuroscientific support, although evidence is more limited. While the hierarchical organization of the visual cortex is well understood^29–32^, the precise manner in which mental imagery is embedded in this representational structure is still a matter of debate^33,34^. Recently, Naselaris and colleagues^5^ showed that visualizing an image leads to the activation of low-level visual features specific to that image within early visual areas V1 and V2, supporting earlier work^2,35^. Some tentative evidence that reactivation within early visual areas correlates with the vividness of mental imagery has also emerged^36^, although the results grouped together the striate and extrastriate cortices, leaving the relation between reinstatement within V1/V2 and vividness unresolved.

In contrast to Hebb’s other two claims, support for his claim that eye movements facilitate neural reactivation during imagery remains largely at the behavioral level. Research indicates that stimulus-specific spatiotemporal fixation patterns elicited during perception are reinstated during retrieval^13–15,37,38^, even in complete darkness^17^. Furthermore, this phenomenon of fixation reinstatement appears to facilitate mental imagery^16,18,38,39^—although, some countervailing evidence exists^15,40,41^. If eye-movements facilitate mental imagery by coordinating shifts of attention to the elements of a remembered visual scene, then it follows that eye-movement reinstatement should be associated with neural reactivation of distributed memory representations. To date, however, there is little neuroscientific evidence supporting this foundational claim of a link between eye movement and imagery.

The goal of the present study was therefore to examine how neural reactivation evoked during mental imagery was related to concurrently measured eye-movement patterns. To capture neural reactivation and eye-movement reinstatement, we collected functional MRI (*f*MRI) and eye tracking data simultaneously while 17 healthy participants viewed and visualized a set of complex colored photographs. In the encoding (perception) condition, participants were repeatedly shown a set of 14 images identified by a unique title and were instructed to remember them in detail. Participants then visualized these images in the retrieval (mental imagery) condition. While this aspect of the experiment is not the focus of the current report, our paradigm was also designed to examine how recency of stimulus presentation influenced neural reactivation patterns. Each retrieval trial began with a sequence of three images (from the set of 14) shown in rapid succession, followed by a cue (title) that identified an image from the set. Participants visualized the image that matched the title, and then rated the vividness of their mental image (Figure 1, In-Scan Task). The recency of the image to be visualized was manipulated in four conditions: long term memory (LTM), wherein the visualized image was not among the three-image sequence; and working memory 1, 2 and 3 (WM1, WM2, WM3), wherein the visualized image was presented in the first, second or third position in the three-image sequence. A post-scan task completed immediately after scanning (Figure 1, Post-Scan Task) served as a behavioral measure of memory acuity. As in the in-scan retrieval condition, participants were shown a sequence of three images (from the in-scan stimulus set) in rapid succession, immediately followed by an image from the set that was either intact or modified (Figure 2). Participants were required to determine whether a subtle change had been made to the image.

**Figure 1.**
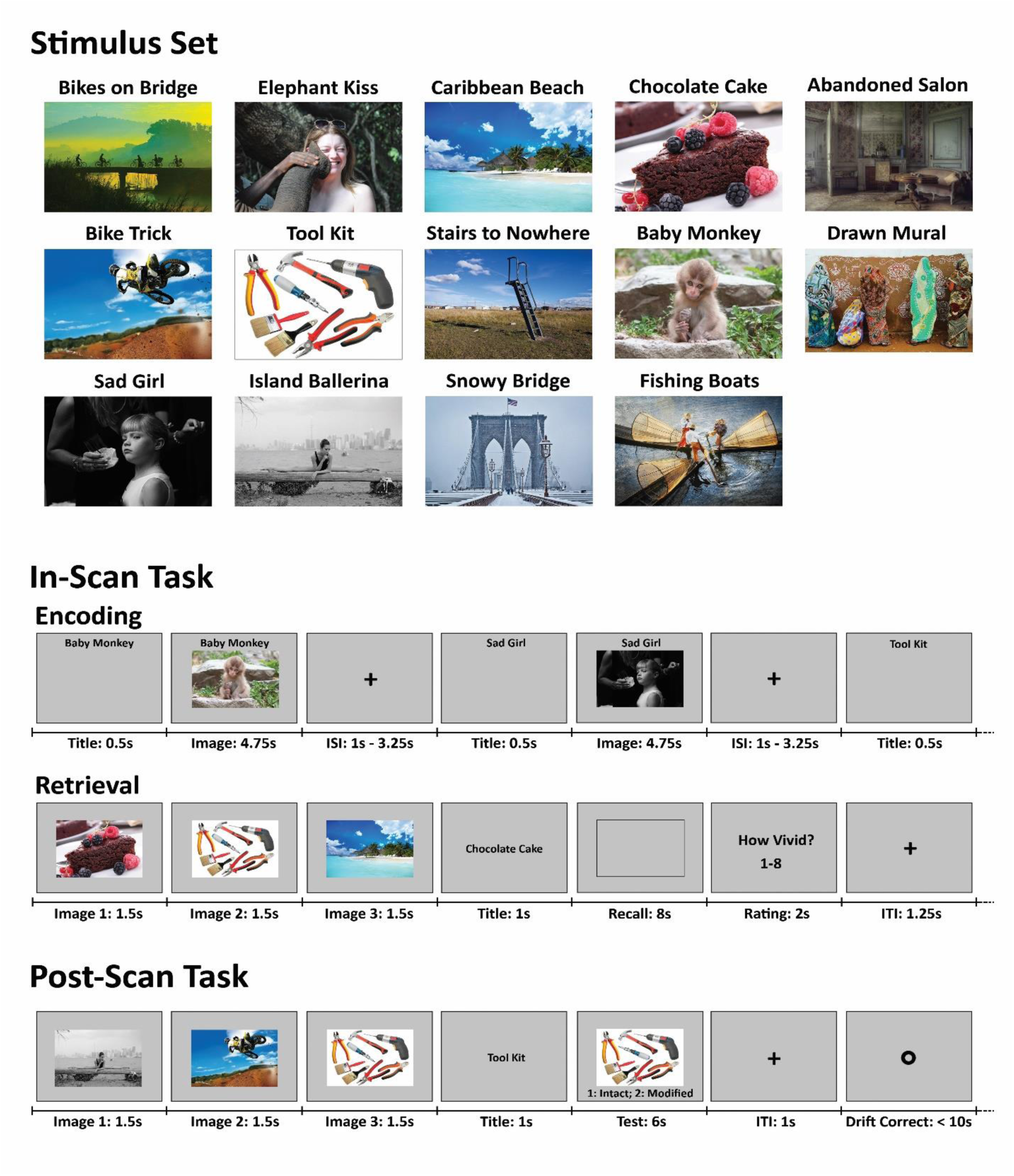
Image Stimuli and Task Procedures. See Methods for an in-depth description of the tasks.

**Figure 2.**
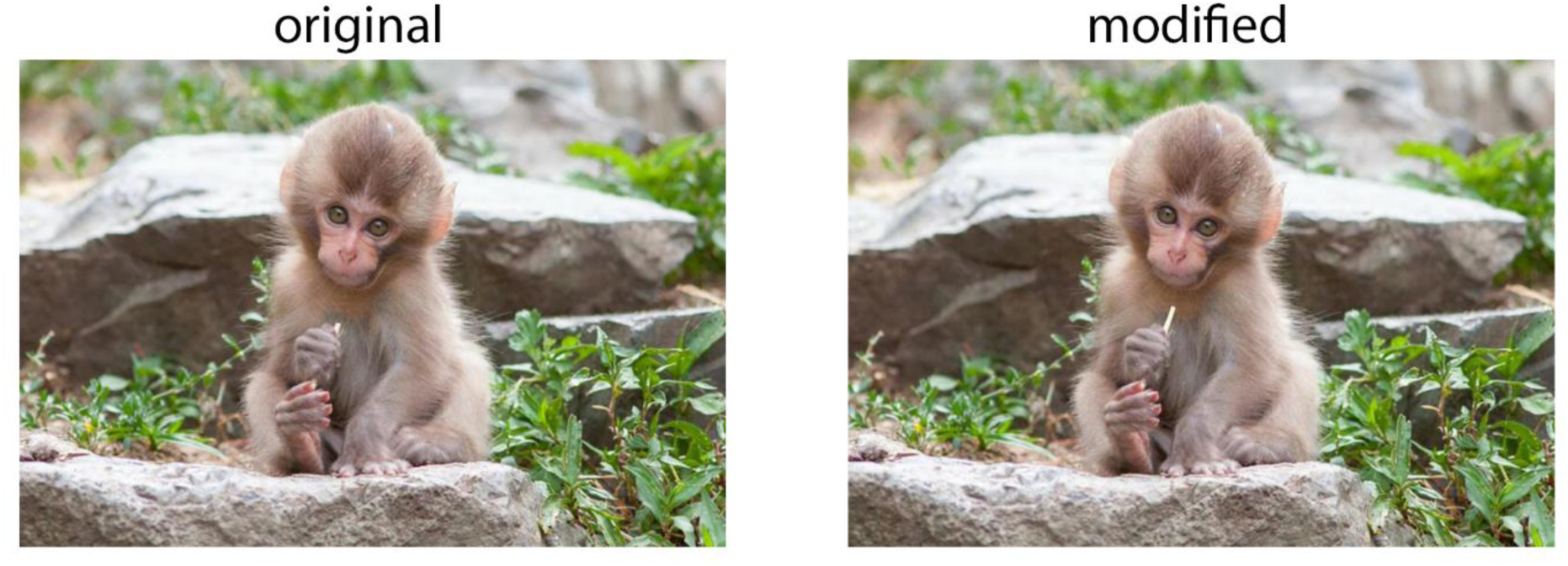
Post-Scan Image Modification. One example of the small modifications that participants were asked to detect during the post-scan behavioral task. Shown images were either modified (right) or identical to the original image (left) held in memory. In this case, the twig the monkey is holding has been lengthened.

We applied MVPA to the *f*MRI signal to quantify the specificity of neural reactivation during mental imagery. We also developed a multivariate spatial similarity analysis method which we applied to the eye tracking data to quantify image-specific patterns of fixation reinstatement. Based on Hebb’s claim that fixation reinstatement should contribute to neural reactivation, we hypothesized that the two metrics should correlate positively, and that the correlation should be strongest at corresponding retrieval time points (i.e. when comparing fixation reinstatement at retrieval-time *x* with neural reinstatement at retrieval-time *x*). Moreover, we hypothesized that individuals capable of conjuring up detailed memories for stimulus items would rely more heavily on eye-movements. If so, we expected post-scan behavioral memory performance to be consistent with in-scan metrics of fixation reinstatement as well as neural reactivation. Finally, we examined Hebb’s claim that reactivation within early visual areas should contribute positively to the vividness of mental imagery. For this, we correlated perceived vividness with neural reactivation in pre-defined visual cortical regions that included the occipital pole and calcarine sulcus.

Our results revealed widespread neural reactivation throughout the time period allocated for visualization. Of interest, imagery vividness ratings correlated positively with reactivation in regions that included the occipital lobe, the ventral and dorsal visual cortex, as well as the calcarine sulcus. Of central importance to our study, neural reactivation was found to correlate positively with fixation reinstatement―even after controlling for neural activity that may have reflected eye movements and fixation position rather than stimulus representations held in memory. The correlation between fixation reinstatement and neural reactivation was strongest when comparing corresponding time points from retrieval trials. To our knowledge, these results provide the first neuroscientific evidence for Hebb’s claim regarding the role of eye movement in mental imagery, as well as support for modern theories of fixation reinstatement, which posit a critical role for eye-movements in the facilitation of memory retrieval^42,43^.

## Results

### Relation between Fixation Reinstatement and Neural Reactivation

We first asked whether patterns of eye fixations made during image encoding are reinstated when the image is visualized at recall. We tested this hypothesis by computing pairwise similarity measures of fixation patterns captured at encoding and recall, which is a form of “representational similarity analysis”^44^ applied to eye-movements. Importantly, participants tend to “contract” their patterns of fixations towards the center of the screen during visualization relative to encoding^14,45,46^. To account for this tendency, we developed a method of spatial fixation pattern alignment based upon the orthogonal Procrustes transform^47,48^. To calculate the measure, two-dimensional fixation density maps were generated for encoding and retrieval trials^49,50^. For each participant, the Procrustes transform was applied, using leave-one-trial-out cross-validation, to spatially align the encoding and retrieval fixation maps. Finally, a trial-specific fixation reinstatement score was calculated by comparing the aligned retrieval trial map’s correlation with the encoding map of the visualized image, relative to that trial map’s average correlation with the remaining 13 encoding maps (see Methods for a detailed description of the measure). Using this novel measure—which greatly outperformed a traditional (unaligned) approach, as measured by recalled image classification accuracy—fixation reinstatement was observed within all recency conditions, with no significant difference between conditions (see Supplementary Figure 1 and Supplementary Table 1).

To assess neural reactivation, we trained a multivariate pattern classifier to discriminate each of the 14 images using brain imaging data from the three runs of encoding task. The trained classifier was then applied to the data from the retrieval task to yield a time point-by-time point estimate of classifier evidence over the course of the visualization window (“cross-decoding”^51,52^). We performed this analysis across the whole brain using feature-selection, which identified voxels concentrated in the posterior half of the brain (Figure 3A). Separate analyses were performed within dorsal, occipital, and ventral cortical regions of interest (ROIs) (Figure 3B and Supplementary Table 5) to separate reactivation in early visual cortex and ventral visual cortex from regions associated with spatial overt attention/eye movements (e.g. intraparietal sulcus) while minimizing multiple comparisons. Neural reactivation was found within the whole-brain and all ROIs for all recency conditions (see Supplementary Figure 2 and Supplementary Table 2).

**Figure 3.**
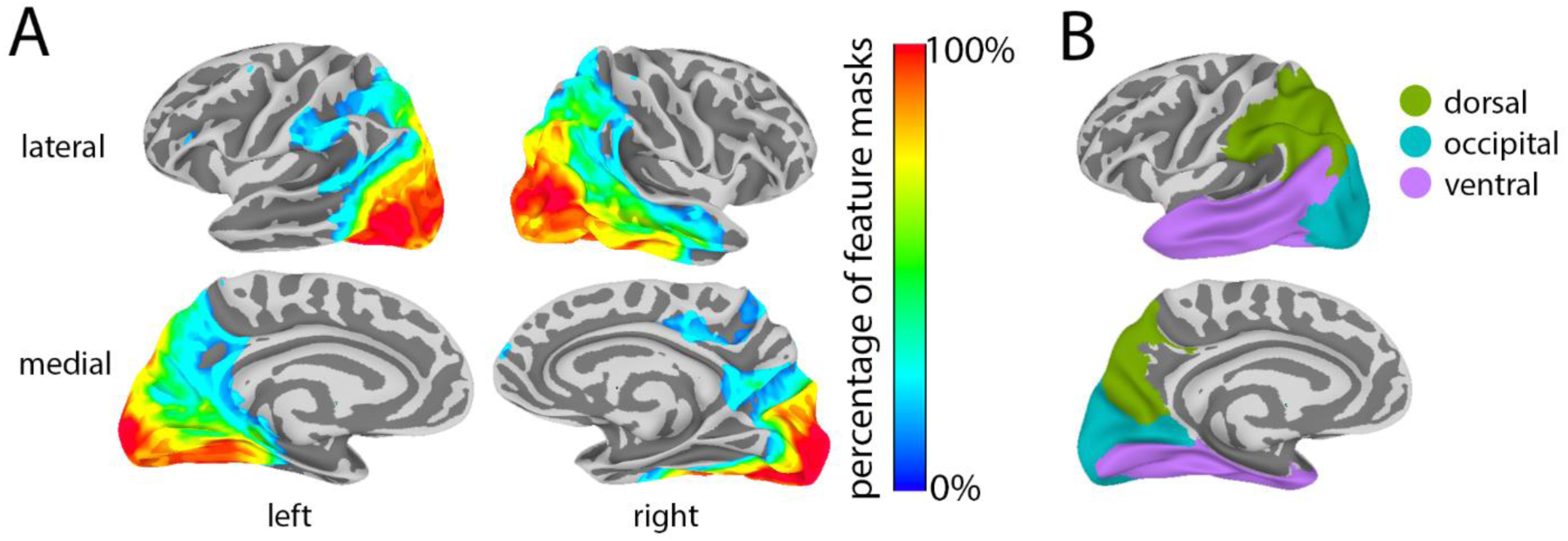
Surface maps of feature and two-stream ROI masks. **A)** Percentage of subject-specific feature masks that contain each voxel. Thresholded at 10%. **B)** Two-stream ROI masks. See Supplementary Table 5 for a list of the FreeSurfer ROIs that compose each region.

Having observed that fixation reinstatement and neural reactivation were both present during our imagery task, we then examined the relationship between the two phenomena. To calculate the correlation between neural reactivation and fixation reinstatement, it was necessary to model several fixed and random factors—including participant, recency condition (LTM, WM1, etc.), recalled image, and recall number (the number of times the current trial’s target image had been previously recalled)—so we used a linear mixed-effects (LME) model. In an analysis of the data from all retrieval trials, we modeled neural reactivation (trial-specific adjusted classifier performance) as a dependent variable (DV), fixation reinstatement (trial-specific fixation reinstatement score) and recall number as scalar independent variables (IV), recency condition as a categorical IV, and participant and image as crossed random effects (random-intercept only, due to model complexity limitations). Statistical assessments were performed using bootstrap analyses.

Figures 4A and 4B illustrate the correlation between fixation reinstatement and neural reactivation. After correcting for multiple comparisons (FDR with one-tailed alpha set to .05), fixation reinstatement correlated positively with reactivation within the feature-selected full-brain when trials from all recency conditions were included (the “all” measure). Correlations specific to recency conditions or limited to signal from specific ROIs were also significant (FDR corrected). We addressed the possibility that the observed correlations were driven by *f*MRI signals caused by similar eye movements made at encoding and retrieval, rather than imagery-related neural patterns *per se*. If true, the similarity between patterns of eye motion made at encoding and at retrieval would result in greater correspondence between patterns of brain activity irrespective of the image being brought to mind (i.e. through random/accidental correlations between eye-movement patterns unrelated to image content). We tested this hypothesis by performing a randomization test for which we generated a null distribution of 1000 randomized “all” correlations (see Methods). For each randomized sample, we randomly reassigned the labels of the visualized images (e.g., all retrieval trials for which “Stairs to Nowhere” was the target image were relabeled as “Chocolate Cake”), and recalculated fixation reinstatement, neural reactivation and their correlation. We found the true “all” correlation to be significantly greater than this null distribution (p = .006), providing strong evidence that the relationship between neural activity and fixations is explained by imagery, and not merely by eye-movement induced patterns unrelated to mental imagery.

**Figure 4.**
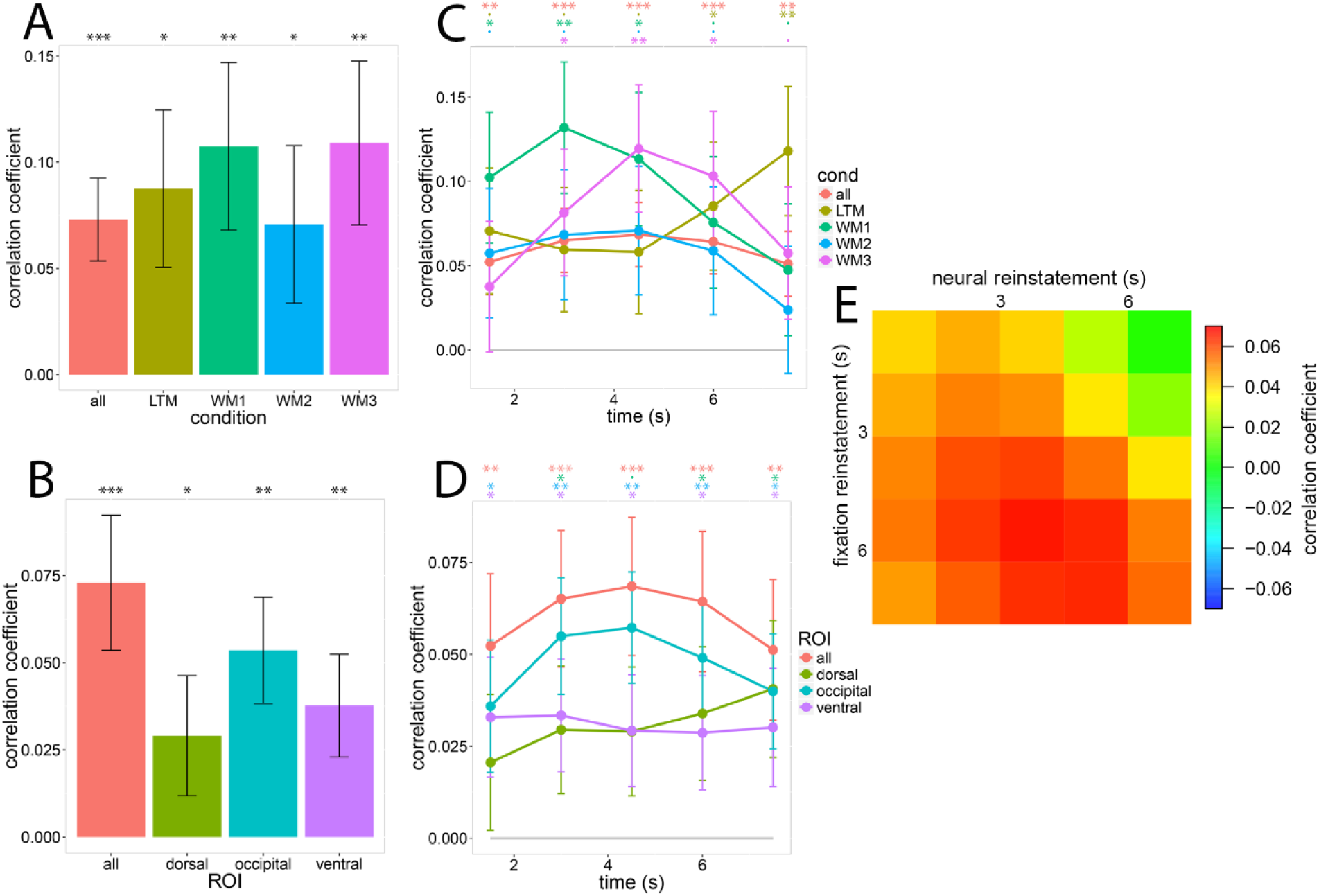
Correlation Between Fixation Reinstatement and Neural Reactivation. Data are represented as correlation coefficient±1 SE; FDR corrected one-tailed p-value: ·<.1, *<.05, **<.01, ***<.001. **A)** The correlation between fixation reinstatement and neural reactivation for each recency condition. The “all” category, which was included in multiple graphs as a point of reference, refers to the full-brain measure that included all recency conditions. **B)** The correlation between fixation reinstatement and neural reactivation for each ROI. **C)** The correlation between fixation reinstatement and neural reactivation for each recency condition divided into retrieval-period temporal windows. **D)** The correlation between fixation reinstatement and neural reactivation for each ROI divided into retrieval-period temporal windows. **E)** The correlation between fixation reinstatement and neural reactivation divided into retrieval-period temporal windows, wherein the columns are neural reactivation windows and the rows are fixation reinstatement windows. See also Supplementary Table 3.

Figures 4C and 4D show the correlation between fixation reinstatement and neural reactivation across the eight-second visualization period. For the feature-selected full-brain correlation including all recency conditions, labeled “all”, the correlation peaked approximately in the middle of the visualization period with all windows significantly greater than zero. Correlations specific to ROIs and recency conditions displayed no consistent temporal pattern, although all groups had at least one significant temporal window—except for ‘WM2’ (FDR corrected). No significant effects were uncovered by a three-way (ROI by recency condition by time) repeated-measures ANOVA performed on the ROI-specific (dorsal, occipital, ventral) correlation data (all *p*s>.30).

Figure 4E shows the relationship between fixation reinstatement and full-brain neural reactivation over time. If eye movements during imagery temporally organize the neural reinstatement of part-images, we hypothesized that the correlation between fixation reinstatement and neural reactivation would be strongest when both measures overlapped in time, i.e. neural reactivation at time *x* should correlate most strongly with fixation reinstatement at time *x*. Qualitatively, the diagonal trend from top-left to bottom-right in figure 4E supports this hypothesis. To test this observation, we first calculated separate correlations between fixation reinstatement and neural reactivation for each time-point combination and each participant. Each correlation was calculated using the LME approach described above, with the exception that participant was not included as a random effect. We then performed an LME analysis with the correlations between fixation reinstatement and neural reactivation as the DV, fixation reinstatement time and neural reactivation time (1-5 scalar valued) as IVs, the absolute difference between fixation reinstatement time and neural reactivation time as an IV, and participant as a random effect. Statistical assessments were performed using bootstrap analyses. We found that the absolute difference between fixation reinstatement time and neural reactivation time correlated negatively with the correlation between fixation reinstatement and neural reactivation (r = −.083, p = .035). In other words, fixation reinstatement and neural reactivation measures were more consistent with each other when taken from time bins that were closer in time, indicating a temporal relationship between the two measures. Overall, the results are consistent with Hebb’s claim that eye movements facilitate the neural reinstatement of part-images during mental imagery.

### Post-Scan Memory Task Performance and Vividness Ratings

The final analyses investigated post-scan memory task performance, vividness ratings and their relation to neural reactivation and fixation reinstatement. Our goal was two-fold: 1) assess whether trials that received high vividness ratings (a subjective measure of imagery) also ranked highly on fixation reinstatement and neural reactivation measures, and 2) determine whether individuals with more detailed memories (those who performed better on the post-scan behavioral memory test) had more specific memory representations (as revealed by in-scan neural reactivation) and relied more heavily on eye-movement recapitulation during imagery.

The post-scan memory task was designed to be difficult, but participants performed above chance, with each individual providing more correct than incorrect answers (% correct: mean = 64.8, p(less than or equal to chance at 50%) < .0001; statistics calculated with bootstrap analyses). To determine whether individuals with good post-scan memory performance (% correct) also obtained high fixation reinstatement and neural reactivation scores, we first computed average fixation reinstatement and neural reactivation scores for each participant, and then we correlated these values with the participants’ memory performance. We covaried out head motion using the maximum displacement (mm) for each subject within the scanner using standard multiple regression. Bootstrap analyses were used to calculate the statistics. Post-scan memory performance correlated strongly with neural reactivation (r = .624, p = .0003, one-tailed), but did not correlate with fixation reinstatement (r = −.015, p = .51, one-tailed).

Given previous findings of a positive correlation between vividness ratings and neural reinstatement^26,36^, we set out to replicate these results, and also to assess whether fixation reinstatement correlated with vividness in the same manner. The within-subject correlations were calculated with a LME model on data from all retrieval trials, wherein either neural reactivation or fixation reinstatement was the dependent variable (DV), Vividness rating and recall number were entered as scalar independent variables (IV), recency condition was a categorical IV, and participant and image were crossed random effects (random-intercept only, due to model complexity limitations). Statistical assessments were performed using bootstrap analyses. Consistent with previous findings, vividness ratings (1-8 scale wherein 1 is very-low and 8 is very-high; mean = 5.57, SD = 1.42) correlated positively with full-brain measures of neural reinstatement (Figure 5A), indicating that image-specific patterns of neural reactivation―an index of memory representation―were more specific during trials perceived as more vivid by the participants. Vividness also correlated with reactivation within the ventral, dorsal and occipital ROIs (Figure 5A and 5C). A two-way (ROI by time) repeated-measures ANOVA revealed that the effects of ROI, time and their interaction were not significant (ROI: F(1.81, 28.90) = .86, p = .42; time: F(2.23, 17.87) = 2.46, p = .13; ROI-time interaction: F(2.04, 32.71) = 2.39, p = .11). Against our hypothesis, no significant positive correlation was observed between vividness and fixation reinstatement (r = .026, p = .10, one-tailed).

**Figure 5.**
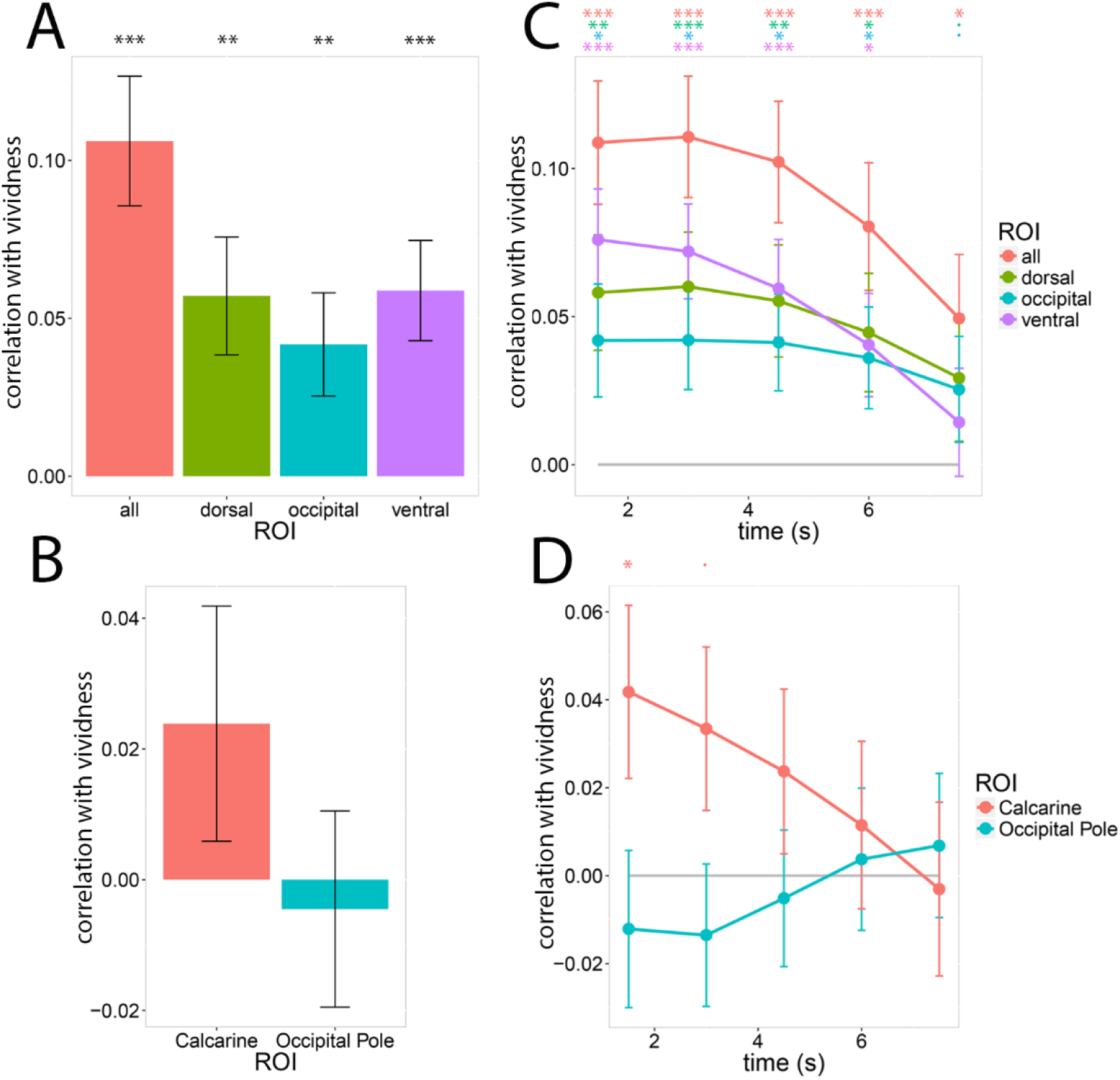
Correlation Between Vividness Rating and Neural Reactivation. Data are represented as correlation coefficient±1 SE; FDR corrected one-tailed p-value: ·<.1, * <.05, **<.01, ***<.001. **A)** The correlation between vividness rating and neural reactivation for each ROI. The “all” category refers to the full-brain measure which included all recency conditions. **B)** The correlation between vividness rating and neural reactivation for the calcarine sulcus and occipital pole. **C)** The correlation between vividness rating and neural reactivation for each ROI divided into retrieval-period temporal windows. **D)** The correlation between vividness rating and neural reactivation for the calcarine sulcus and occipital pole divided into retrieval-period temporal windows. FDR multiple comparison correction was applied sequentially, starting at the first time-point. See also Supplementary Table 4.

We also tested Hebb’s claim^12^ that neural reactivation in early visual areas elicited more vivid visual mental imagery. Looking specifically at the signal from early visual ROIs, namely the occipital pole and calcarine sulcus, we found no significant correlations between reactivation and vividness after FDR correction (Figure 5B and 5D). A two-way (ROI by time) repeated-measures ANOVA revealed that the effects of ROI, time and their interaction were non-significant (ROI: F(1, 16) = 1.90, p = .18; time: F(1.46, 23.37) = 1.25, p = .29; ROI by time interaction: F(1.33, 21.33) = 2.00, p = .16). Because neural reinstatement decreased approximately linearly over retrieval time (Supplementary Figure 2), an ANOVA—which does not assume any relation between time points—may be underpowered. To address this issue, we ran an LME model that assumed a linear relation between time points. In this model, the correlation between vividness and neural reinstatement, calculated for each subject-ROI-time combination, was the DV; ROI, time and their interaction were IVs; and participant was a random effect. The main effect of ROI and the ROI-time interaction were significant, indicating that the correlation between neural reinstatement and vividness was significantly stronger within the calcarine sulcus than the occipital pole—particularly near the start of the visualization period (ROI: coefficient = .183, p = .0058; time: coefficient = −.109, p = .09; ROI-time interaction: coefficient = −.146, p = .024; calculated via bootstrap analyses). Based upon this finding, we re-analyzed the correlation between vividness and neural reactivation over time by assessing each time window sequentially, starting from the beginning of the visualization period and including all previous time windows in a multiple comparison analysis using FDR. Using this method, we found the first (0-1.5 sec) visualization time window for the calcarine sulcus to be significant (0-1.5 sec: r = .041, p = .03, one-tailed), whereas all other windows for both ROIs were not significant.

To determine whether these results were limited by our ability to detect neural reactivation in these early visual regions, we assessed neural reactivation over time within the occipital pole and calcarine sulcus. We performed random effects (subjects and items) bootstrap analyses for each retrieval time point—controlling for multiple comparisons by assessing the time points sequentially using FDR, as described above. Only the first visualization time window (0-1.5 sec) was found to be significant for the calcarine sulcus (0-1.5 sec: adjusted classifier confidence (%) = 1.51, p = .03, one-tailed), mirroring the correlation results.

These results document the spatiotemporal relationship between neural reactivation and the perceived vividness of mental images. While we observed significant correlations between vividness and reactivation across the visual cortex, we found limited evidence in support of Hebb’s claim12 that reactivation in early visual cortices leads to vivid mental imagery. That being said, our capacity to detect reactivation in early visual cortices may have been affected by our study design, which allowed subjects to move their eyes during visualization, a limitation that we address further in the discussion.

## Discussion

The primary goal of the current study was to test whether eye movements contribute to the creation of mental images by examining the relationship between fixation reinstatement—as measured by a novel Procrustes-transform-based algorithm—and neural reactivation. Our results provide significant evidence in favor of Hebb’s claim^12^ that eye movements help coordinate the construction of mental images. We observed a significant positive correlation between a novel measure of fixation reinstatement that accounts for the contraction of fixations during imagery, and neural reactivation. This correlation increased when fixation reinstatement and neural reactivation metrics were calculated for time points that were closer in time, demonstrating that the two phenomena peaked in synchrony, and establishing a link between eye movement and the neural mechanism of mental imagery.

Previous research has only assessed the link between fixation reinstatement and mental imagery using behavioral measures of imagery rather than neural reactivation. For example, Laeng and Teodorescu^16^, and Johansson et al.^18^, found that the degree of fixation reinstatement predicted behavioral performance on an imagery task. Thus, our findings provide the first direct neuroimaging evidence for Hebb’s claim and the currently dominant fixation reinstatement theories^14,40,42,43^.

Our analyses also addressed the relationship between fixation reinstatement, neural reactivation and behavioral memory performance. Based on findings linking fixation reinstatement^16,18^ and neural reactivation^6,7,24^ to memory performance, we predicted that both in-scan fixation reinstatement and neural reactivation would correlate with performance on the post-scan memory task. We also predicted similar patterns of correlations with in-scan ratings of imagery vividness. Our results were partially congruent with these predictions. We observed that neural reactivation correlated strongly with both objective and subjective behavioral measures of memory performance, but that fixation reinstatement was a poor predictor of either form of behavior.

Research into the relationship between fixation reinstatement and memory acuity has been mixed^53,15,40,16,18,38^. For example, when fixations were constrained to a region that either did or did not correspond to the previous location of objects to be recalled, Johansson and Johansson^39^ found that memory performance was superior in the “corresponding” condition, whereas Martarelli and Mast^41^ did not. These inconsistent results may be due to differences in the features to be recalled: spatial features (orientation and relative position) in Johansson and Johansson^39^, and primarily non-spatial features (e.g. color) in Martarelli and Mast^41^. Consistent with this interpretation, de Vito et al.^54^ demonstrated that incongruent eye movements preferentially disrupt spatial recollection. Our objective measure of memory performance, the post-scan change detection task, included both spatial (e.g. size, position) and non-spatial (e.g. color, object identity) image modifications. Therefore, we may have observed a larger correlation between in-scan fixation reinstatement and post-scan change detection if the task only had spatial modifications. Similarly, subjective vividness ratings reflected an overall impression of the crispness of the mental image, rather than its spatial features.

In summary, we provided the first evidence that fixation reinstatement is linked to neural reactivation, thereby supporting one of the pillars of Hebb’s theory of imagery, as well as current fixation reinstatement theories which also posit that reciprocal facilitation occurs between fixation reinstatement and internal memory representations^42,43^. Nonetheless, we must consider alternative interpretations of our results. It is possible that the observed correlation between fixation reinstatement and neural reactivation was predominantly driven by neural signals that reflected eye position, rather than the reactivation of a mental image. A randomization test designed to address this possibility provided strong evidence against this hypothesis. Moreover, positive correlations based on signal limited to the occipital and ventral ROIs provide further evidence against this hypothesis, as these ROIs exclude areas strongly associated with eye movement control, such as the frontal eye fields and posterior intraparietal sulcus^55,56^.

Finally, while our correlational findings reveal a relationship between eye movement and imagery, they cannot conclusively determine the causality and directionality of this relationship. To directly address this unresolved issue, future research could take advantage of the high temporal resolution of techniques such as magnetoencephalography to link distinct patterns of neural activity to specific portions of seen and imagined complex images. If reciprocal facilitation occurs between fixation reinstatement and internal memory representations^42,43^, then neural reactivation should predict, and be predicted by, eye movements towards the location associated with the neural activity pattern.

We also tested Hebb’s claim^12^ that highly vivid mental imagery requires cortical reactivation within early visual areas, i.e. V1 and V2. As such, we hypothesized that reactivation within the occipital pole and the calcarine sulcus would correlate positively with vividness. Consistent with previous findings^26,27,36^, we observed correlations between vividness and reactivation within dorsal, ventral and occipital ROIs that were sustained throughout the retrieval trial. Looking specifically at early visual areas, a significant correlation was observed between vividness ratings and reinstatement within the calcarine sulcus (the brain region wherein V1 is concentrated^57^), but not the occipital pole, in the first 1.5 seconds of visualization.

The simplest explanation for the null result within the occipital pole is that vivid mental images can be conjured up without its contribution to neural reactivation. However, other factors need to be considered. First, St-Laurent, Abdi, and Buchsbaum^26^ found that activity levels within the occipital pole correlated strongly and positively with the perceived vividness of videos mentally replayed from memory, which suggests that this area contributes to the perceived vividness of mental imagery. Second, we only observed evidence of neural reactivation during the first 1.5 seconds of visualization within the calcarine sulcus, and not within the occipital pole, mirroring our correlation results. This finding suggests that the observed correlation between reactivation within early visual regions and vividness was limited by our ability to detect reactivation within these regions.

Research by Naselaris et al.^5^ provides strong evidence of reactivation of neural patterns associated with low-level visual features within the early visual cortex during scene imagery. One significant methodological difference between this study and our own is that the authors asked their participants to fixate centrally throughout their task, thereby eliminating the natural eye-movements that occur during mental imagery—which were the explicit focus of our study. This significant constraint on the participants’ fixations would have eliminated the variance caused by the image’s neural representation shifting across the retinotopically-organized early visual cortex due to eye movements, but at the cost of being able to study the functional role of eye-movements during imagery^18^. Note that the occipital pole and posterior calcarine sulcus are predominantly responsible for central vision, which has high spatial resolution, whereas the anterior calcarine sulcus is predominantly responsible for peripheral vision, which has relativity low spatial resolution^57^. Consequently, visual representations within the calcarine sulcus should be less sensitive to eye movements than visual representations within the occipital pole, which is consistent with our results. We therefore suspect that free eye-movements may have caused our null reactivation finding within the occipital pole. By extension, our methods could not adequately quantify the correlation between vividness and reactivation within the occipital pole and calcarine sulcus, and limited our capacity to test Hebb’s claim that early visual cortical reactivation leads to vivid imagery. To preserve ecological validity, future research concerning neural reactivation during mental imagery should avoid artificial constraints on fixations, and instead develop and utilize measures of neural reactivation that explicitly model the effect of eye movements on neural activity within the visual cortex.

## Conclusion

In conclusion, the results from this study support the three major claims of the Hebbian theory of mental imagery: 1) imagery involves the reinstatement of perceptual neural activity; 2) reinstatement of fixations during imagery facilitates neural reinstatement; 3) the vividness of mental imagery is associated with reactivation within early visual areas (calcarine sulcus). The findings reported here provide a promising avenue to establish how fixations contribute to the neural processes underlying mental imagery. Future work should clarify the fine-scale temporal relationship between eye-movement reinstatement and memory reactivation in a way that can unravel the causal connection between these interacting neural processes.

## Methods

### Participants

Twenty-three right-handed young adults (6 males and 17 females, 20-30 years old [mean: 24.1], 14-21 years of education [mean: 16.9]) with normal or corrected-to-normal vision and no history of neurological or psychiatric disease were recruited through the Baycrest subject pool, tested and paid for their participation per a protocol approved by the Rotman Research Institute’s Ethics Board. Subjects were either native or fluent English speakers and had no contraindications for MRI. Data from six of these participants were excluded from the final analyses for the following reasons: excessive head motion (2), poor eye tracking signal (1), misunderstood instructions (1), fell asleep (2). Thus, seventeen participants were included in the final analysis (5 males and 12 females, 20-28 years old [mean: 23.8], 15-21 years of education [mean: 17.1]).

### Stimuli

Nineteen complex colored photographs were gathered from online sources and resized to 757 by 522 pixels in Adobe Photoshop. Five images were used for practice, and the remaining 14 were used during the in-scan and post-scan tasks (Figure 1). Each image was paired with a short descriptive title in 30-point Courier New font during in-scan encoding; this title served as a retrieval cue during the in-scan and post-scan memory tasks. Four different “modified” versions of each image were also created using Adobe Photoshop for a post-scan memory test: a minor local element of the image was either added, removed or transformed in a way that was realistic and congruent with the image (Figure 2).

## Procedure

### In-Scan

Before undergoing MRI, participants were trained on a practice version of the task incorporating five practice images. Inside the scanner, participants completed three encoding runs and six retrieval runs of functional MRI. To keep participants engaged with the task, we interspaced the encoding and the retrieval runs (each encoding run was followed by two retrieval runs). A high-resolution structural scan was acquired between the 6th (retrieval) and 7th (encoding) functional runs, which provided a mid-task break. Eye-tracking data was acquired during all functional runs.

Encoding runs were 7m 18s long. Each run started with 10s of warm up during which instructions were displayed on-screen. Each trial began with a title shown in the top portion of the screen (0.5s; font = Courier New, font size = 30), followed by the appearance of the matching image in the center of the screen (4.75s; the title remained visible above the image). Images occupied 757 by 522 pixels of a 1024 by 768 pixel screen. Between trials, a cross-hair appeared in the center of the screen (font size = 50) for either 1s, 1.75s, 2.5s or 3.25s.

Participants were instructed to pay attention to each image and to encode as many details as possible so that they could visualize the images as precisely as possible during the retrieval task. During the second and third encoding runs, participants were encouraged to pick up details they had missed and to integrate them into their memory representation. Each image was shown four times per run, for a total of 12 encoding trials per image throughout the experiment. Within each run, the entire set of images was shown in a randomized order before the set could be shown again (e.g. each image needed to be shown twice before an image could be presented for the third time).

Retrieval runs were 8m 17s long, starting with 13 seconds of warm up during which instructions appeared on-screen. Each trial began with three 757 by 522 images shown in succession in the center of the screen for 1.5s each. Then, an image title appeared in the center of the screen for 1s (font = Courier New, font size = 30). For most trials, this title matched one of the three images in the sequence. The first, second and third image from the sequence were each cued during working memory conditions 1, 2 and 3, respectively (WM1, WM2, and WM3). WM1, WM2 and WM3 trials each corresponded to 1/4 of the total number of trials. In the remaining 1/4 of trials, the title corresponded to an image from the stimulus set that was not included in the sequence (the long-term memory condition, LTM). After 1s, the title was replaced by an empty rectangular box shown in the center of the screen (8s), and whose edges corresponded to the edges of the stimulus images (757 by 522 pixels). Participants were instructed to visualize the image that corresponded to the title as accurately and in as much detail as they could within the confines of the box. Once the box disappeared, participants were prompted to rate the vividness of their mental image on a 1-8 scale (2s) using two four-button fiber optic response boxes (one in each hand; 1 = left little finger; 8 = right little finger). Between each trial, a cross-hair (font size = 50) appeared in the center of the screen for 1.25s. Participants were instructed to attribute ratings of 4 or 5 for trials whose vividness felt “average for them”. There were 28 trials per run (seven trials in each condition: WM1, WM2, WM3 and LTM), and 42 trials per condition for the entire scan.

### Post-Scan

A post-scan test was conducted shortly after scanning to obtain behavioral measures of memory specificity as a function of task condition for the same 14 images encoded and retrieved inside the scanner. For each original image, four modified versions were created (Figure 2) which were used as difficult recognition probes to test each individual’s memory acuity for the 14 images. Participants were instructed on the new task and completed a practice that included the five practice images shown during pre-scan training. The task involved four consecutive retrieval blocks separated by short breaks and, if needed, eye-tracking recalibration. For each trial, three images (757 by 522 pixels) from the set were presented consecutively in the center of a 1024 by 768 pixel screen for 1.5s each. Then, in a manner analogous to the in-scan retrieval task, an image title appeared in the center of the screen (1s; font = Courier New, font size = 30) that either matched the first (WM1), second (WM2) or third (WM3) image from the sequence, or that corresponded to an image from the set that was not included in the sequence (LTM; 1/4 of trials were assigned to each condition). The title was followed immediately by a version of the corresponding image that was either intact or modified. Participants were given 6s to determine whether the image was intact or modified using a keyboard button press (right hand; 1 = intact, 2 = modified). After 6s, the image was replaced by a 1s fixation cross (font size = 50) during which participants’ response could still be recorded. The images shown in the 3-image sequence were always intact. Each of the four modified versions of an image appeared only once in the experiment (for a single trial), each in a different condition. During the inter-trial interval, participants were required to fixate on the inner portion of a small circle in the center of the screen. The experimenter pressed a button to correct for drifts in calibration and to trigger the onset of the next trial. Participants were informed they could move their gaze freely during the rest of the trial.

For each original image, the four modified versions (Figure 2) were arbitrarily labeled modified images 1 to 4. Across participants, we counterbalanced the conditions in which an image was tested within each block, the condition to which an image’s modified version was attributed, and the block in which a modified image’s version appeared.

### Setup and Data Acquisition

Participants were scanned with a 3.0-T Siemens MAGNETOM Trio MRI scanner using a 12-channel head coil system. A high-resolution gradient-echo multi-slice T1-weighted scan coplanar with the echo-planar imaging scans (EPIs) was first acquired for localization. Functional images were acquired using a two-shot gradient-echo T2*-weighted EPI sequence sensitive to BOLD contrast (22.5 × 22.5 cm field of view with a 96 × 96 matrix size, resulting in an in-plane resolution of 2.35 × 2.35 mm for each of 26 3.5-mm axial slices with a 0.5-mm interslice gap; repetition time = 1.5 sec; echo time = 27ms; flip angle = 62 degrees). A high-resolution whole-brain magnetization prepared rapid gradient echo (MP-RAGE) 3-D T1 weighted scan (160 slices of 1mm thickness, 19.2 × 25.6 cm field of view) was also acquired for anatomical localization.

Both the in-scan and the post-scan task were programmed with Experiment Builder version 1.10.1025 (SR Research Ltd., Mississauga, Ontario, Canada). In the scanner, stimuli and button press responses were presented and recorded using EyeLink 1000 (SR Research Ltd., Mississauga, Ontario, Canada). Visual stimuli were projected onto a screen behind the scanner made visible to the participant through a mirror mounted on the head coil. In-scan monocular eye movements were recorded with an EyeLink 1000 infrared video-graphic camera equipped with a telephoto lens (sampling rate 1000Hz) set up inside the scanner bore behind the participant’s head. The camera picked up the pupil and corneal reflection from the right eye viewed from the flat surface mirror attached inside the radio frequency coil. Nine-point eye movement calibration was performed immediately before the first functional run. If needed, manual drift correction was performed mid-scan immediately prior to the onset of the next trial, and calibration was re-done in-between subsequent runs.

Post-scan stimuli were presented on a 19-in. Dell M991 monitor (resolution 1024×768) from a 24-inch distance. Monocular eye movements (the most accurate eye was selected during calibration) were recorded with a head-mounted Eyelink II eye tracker (sample rate 500 Hz) set to detect the pupil only. Eye movement calibration was performed at the beginning of the experiment, and drift correction (>5°), if needed, was performed immediately prior to the onset of each trial.

In-scan and post-scan eye tracking and behavioral data (vividness ratings, accuracy, and response time) were analyzed with Dataviewer version 1.11.1 (SR Research Ltd.). Saccades were determined using the built-in EyeLink saccade-detector heuristic. Acceleration (9500°/s/s) and velocity (30°/sec) thresholds were set to detect saccades greater than 0.5° of visual angle. Blinks were defined as periods in which the saccade-detector signal was missing for three or more samples in a sequence. Fixations were defined as the samples remaining after the categorization of saccades and blinks.

For the post-scan memory task, regions of interest (ROIs) were defined manually a priori for each image. A rectangular shape was drawn over each area of the image where a modification was introduced during the change-detection task, totaling four ROIs per image. Variations in the shape and orientation of these rectangles was dictated by the nature of the change, but strict counterbalancing insured that each variation was assigned to different conditions in a non-biased manner across participants.

### *f*MRI and Neural Reactivation Measures

All statistical analyses were first conducted on realigned functional images in native EPI space. Functional images were converted into NIFTI-1 format, motion-corrected and realigned to the average image of the first run with AFNI’s^58^ *3dvolreg* program, and smoothed with a 4-mm FWHM Gaussian kernel. The maximum displacement for each EPI image relative to the reference image was recorded.

For each subject, shrinkage discriminant analysis^59^ (SDA) was used to train a pattern classifier to discriminate between the set of 14 images using fMRI data from the encoding runs. The full-brain, “all” ROI, pattern classifier was trained in two steps. First, a multivariate searchlight analysis using an 8mm radius and using leave-one-run-out cross-validation was used to detect regions with above chance performance in classifying the label associated with the 14 images. The searchlight classification accuracy maps were then thresholded at Z>1.65 (binomial distribution with chance accuracy = 1/14) to create separate feature masks for each subject (Figure 3A). A second SDA classifier was then trained on the encoding runs using all voxels falling inside the subject’s feature mask, producing a final full-brain classifier that could be used to evaluate image-specific reactivation during the retrieval task.

For the ROI reinstatement analyses, the subject-specific feature masks (Figure 3A) were divided into “dorsal”, “occipital” and “ventral” regions (Figure 3B), based upon Two-Streams hypothesis^60^—where “occipital” ROIs are not predominantly associated with one of the streams (see Supplementary Table 5 for a list of the FreeSurfer ROIs that compose each region). Three SDA classifiers per subject, one for each ROI, were then trained on the encoding runs using all voxels falling inside the subject’s feature mask and the ROI’s mask. In a similar manner, occipital pole and calcarine sulcus ROI analyses were performed with two SDA classifiers per subject, one for each ROI, but they were trained using all voxels within the corresponding FreeSurfer bilateral ROIs.

The SDA pattern classifiers trained on the set of encoding trials were then applied to data from the same brain regions acquired during the retrieval task. First, the time-series data for each individual memory trial was divided into 16 intervals of 1.5 seconds (spanning 0-24 s), where the first interval (0-1.5 s) is aligned to the start of the trial, which is defined as the onset of the first image from the three-image sequence (see Figure 1). Next, the SDA classifiers were applied to each time-point of each retrieval trial, producing a time-course of classifier confidence for each trial. To control for the cortical activation caused by the recency condition (i.e. the three images shown at the onset of retrieval trials), we produced an “adjusted classifier confidence” (see Supplementary Figure 3 for an explanatory diagram). A trial’s classifier confidence was adjusted by subtracting the average classifier confidence for trials during which the target trial’s visualized image was shown in the same serial position (or not shown at all, as in the “LTM” condition), but was *not* retrieved (e.g. for a “WM2” trial where the visualized image “Baby Monkey” is shown in position 2, the average classifier confidence for all “non-WM2” trials where “Baby Monkey” is shown in position 2 is subtracted). With this metric, a value greater than 0 indicated neural reinstatement. The adjusted classifier confidence for each time-point was smoothed by convolving the data with a Gaussian filter (SD = 2 seconds), and a single adjusted classifier confidence score was calculated for each trial by averaging across the five time-points corresponding to the visualization period (5.5-13 seconds offset by 6 seconds, i.e. 11.5-19s, to account for hemodynamic delay; the last 0.5 seconds were cut to avoid overlap with the vividness judgment).

### Fixation Reinstatement Measure

Fixation reinstatement―the similarity between spatial fixation patterns during encoding and retrieval―was assessed by calculating the correlation between fixation density maps^49,50^. To create fixation density maps, a 3D Gaussian distribution was centered on each fixation made during the trial. The Gaussian’s "height" was proportional to the fixation’s duration and its width was such that one standard deviation was about 1 degree of visual angle, approximating the width of the fovea. For each pixel on the screen, the different Gaussians’ values (one per fixation) at that pixel were summed, and the resulting map was normalized so that the sum over all pixel values was 1. To speed up computational processing time, maps were calculated at 1/32 the resolution of the original screen.

Multiple studies have shown that the dispersion of fixations is lower when an image is visualized rather than perceived. This effect varies significantly between individuals^14,17,45^ and is linked to differences in spatial imagery, so that participants with higher OSIVQ scores^61^ have more spatially constrained fixations^46^. Counter-intuitively, those with superior spatial imagery may therefore show less similarity between encoding and retrieval fixation density maps. To control for this contraction of fixations during mental imagery, we aligned encoding and retrieval fixation density maps using the orthogonal Procrustes transformation^47,48^—a geometric transformation that uses translation, rotation and scaling to jointly minimize the distance between a two sets of paired points.

To calculate fixation similarity, we first generated encoding fixation maps by combining fixations made within the spatial boundaries of the image for the entire period when it was on screen. Encoding fixations were combined across trials for each subject-image combination (14 encoding maps per subject). At retrieval, fixations were divided into 29 time windows that spanned the trial’s visualization period (window duration/width = 1s, temporal distance between windows/stride = 0.25s), Fixations that straddled the border of a window had their durations limited to the duration spent within the window. Retrieval maps were created by pooling all fixations made within the on-screen rectangular visualization box, for each subject-image-time window combination (fixations were pooled across trials; 14*29 maps per subject). Trial-specific retrieval maps were also generated for each subject-trial-time window combination (29 maps per trial per subject). For cross-validation, we also generated retrieval maps for each subject-trial-time window combination that incorporated all fixations made by a subject within a certain time window during trials with the same target image as a particular trial (across conditions)— excluding that trial’s fixations.

To correct for each subject’s individual tendency to systematically alter fixations at retrieval, retrieval maps were aligned with encoding maps using the Procrustes transformation. Crucially, alignment parameters were calculated in a single step using a subject’s encoding and retrieval fixation data from all 14 stimulus images, yielding a transformation matrix that optimally rotates the set of retrieval fixation maps to match the set of encoding fixation maps. Thus, this method does not compute a separate transformation for each *image*, but rather discovers a single transformation that optimally aligns the two *sets* of 14 fixation maps. Moreover, to evaluate the test performance of the Procrustes method, a leave-one-out cross-validation approach was used in which the transform was calculated on all trials except for the “left out” test trial. Specifically, a separate Procrustes transformation matrix was calculated for each subject-trial-time window combination by jointly aligning two 14 by 768 matrices―one for encoding and one for retrieval. Matrix rows represented the 14 stimulus images, and columns represented pixel-specific elements from the vectorized fixation maps. Rows from a subject’s encoding matrix corresponded to vectorized encoding fixation maps (one map per image, with fixations combined over trials). A different retrieval matrix was created for each subject-trial-time window combination: elements from the target image’s row corresponded to the “cross-validation” fixation map that excluded that trial’s fixations but included fixations from other trials with the same target image made within the target time window. Other rows corresponded to the other images’ vectorized retrieval fixation maps (combined across trials) for the target time window.

For each subject-trial-time window combination, alignment resulted in a transformation matrix that was used to transform the fixation map specific to that retrieval trial and time window. The transformed retrieval fixation map was then correlated with each of the subject’s 14 encoding fixation maps (one for each image). To match the temporal profile of our neural reactivation measure, correlations for each of the 29 time windows were reduced to 5 values by convolving them with 5 Gaussians (means = 0.8, 2.4, 4.0, 5.6, 7.2 sec; SD = 2 sec); a single non-temporal correlation value was also calculated as the mean of the 5 temporal values. For both the 5 temporal correlations and the non-temporal correlation, the final fixation reinstatement value was calculated as the difference (subtraction) between the correlation with the encoding fixation map corresponding to that trial’s target image, and the average correlation with the other fixation maps for the non-target images. For this measure, a value greater than zero indicates fixation reinstatement.

### Bootstrap and Randomization Statistics

All bootstrap statistics were calculated with 10000 samples. For the calculation of correlation statistics using a linear mixed effects (LME) model, bootstrap analyses were calculated with the BootMer function^62^. For the calculation of mean statistics using a LME model, an array was created with each dimension representing a random effect—in this case, participants (17 rows) and images (14 columns). Each element of the array is the mean value for the element’s combination of random effects (e.g. row 3, column 5 contains the mean value for participant 3, image 5). To generate a bootstrap distribution of the mean, 10000 new matrices were generated by randomly sampling the rows and columns of the original matrix with replacement, and then the 10000 means of the matrices’ elements were calculated. For the paired-samples variant of the preceding procedure, each element of the array was a difference of means (i.e. the difference between the means generated by two different fixation similarity algorithms).

To address the possibility that the observed correlation between fixation reinstatement and neural reactivation was the result of *f*MRI signals directly caused by eye movements, as opposed to imagery-related neural activity patterns, we performed a randomization test. If the null hypothesis is true, then similarity between patterns of eye-movements made at encoding and at retrieval would result in greater correspondence between patterns of brain activity *irrespective of the image being brought to mind*. If so, then the similarity of retrieval fixation patterns made while visualizing image *x* to encoding fixation patterns made while perceiving image *y* should predict neural reactivation of image *y* to the same degree that fixation reinstatement of image *x* predicts neural reactivation of image *x*. We generated a null distribution by randomly reassigning the labels of the target images visualized during retrieval trials, and then re-calculating the correlation between fixation reinstatement and neural reactivation. Specifically, each retrieved image was randomly reassigned to another image from the set with two constraints: an image was never assigned to itself, and an image was never assigned to more than one image (i.e., all trials during which “Baby Monkey” was the retrieved image were assigned the same new label). After image assignment, all variables that were dependent on the identity of the retrieved image were recalculated (i.e. recency condition, fixation reinstatement, and neural reactivation), and the correlation between fixation reinstatement and neural reactivation was stored. This process was repeated 1000 times, producing a 1000 sample null distribution, which was then compared to the original correlation between fixation reinstatement and neural reactivation.

## Author Contributions

Conceptualization, M.B.B., B.R.B., M.S.; Methodology, M.B.B., B.R.B., M.S.; Software, M.B.B., B.R.B.; Formal Analysis, M.B.B., B.R.B., M.S.; Investigation, M.S., C.D., D.A.M.; Resources, C.D.; Data Curation, M.B.B., B.R.B.; Writing – Original Draft, M.B.B., M.S., B.R.B.; Writing – Review and Editing, M.B.B., B.R.B., M.S., J.D.R., D.A.M.; Visualization, M.B.B., B.R.B., M.S.; Supervision, B.R.B., J.D.R.

## Acknowledgements

We thank Morris Moscovitch and Jordana Wynn for their insightful discussions. This work was supported by a NSERC Discovery Award. There are no conflicts of interest.

